# Large-scale exposure of southern seabirds to avian cholera *Pasteurella multocida*: scavengers as sentinels at different spatial scales

**DOI:** 10.64898/2026.10.12.758445

**Authors:** Jeremy Tornos, Amandine Gamble, Augustin Clessin, Tristan Bralet, Mathilde Lejeune, Maëlle Connan, Peter G. Ryan, Henri Weimerskirch, Jacob Gonzalez-Solis, Paulo Catry, Christophe Barbraud, Karine Delord, Vincent A. Viblanc, Aline Flechet, Juliette Baron, France Caza, Amira Tounsi, Sophia Ferchiou, Samuel Perret, Charles-André Bost, Antje Steinfurth, Rozenn Le Net, Jean-Baptiste Thiebot, Fabrice Lebouard, Célia Lesage, Yves St-Pierre, Hubert Gantelet, Thierry Boulinier

**Author notes:** Corresponding author: Thierry Boulinier CEFE UMR 5175 CNRS, Université Montpellier, EPHE, IRD 1919 Route de Mende, 34293 Montpellier, France.

## Abstract

Seabird communities represent spatially structured social networks within which potential sentinels could inform us where multi-host pathogens could be present. *Pasteurella multocida,* the agent of avian cholera, is responsible for recurrent massive seabird mortality events on remote Amsterdam Island, Southern Indian Ocean, but little is known about its broad spatial occurrence. To explore its distribution and whether avian scavengers and predators could be used as sentinels, we screened 18 southern seabird species from 11 islands for antibodies against *P. multocida* (2634 plasma samples). We detected widespread but heterogeneous exposure among host populations. High seroprevalences were found in widely roaming predators and scavengers (giant petrels (*Macronectes* spp), and brown skua (*Stercorarius antarcticus)*), which could represent efficient sentinels at broad scales. Antibodies were also detected over a series of years in a non-migratory scavenger, the lesser sheathbill (*Chionis minor*) and an abundant penguin species, the king penguin (*Aptenodytes patagonicus)*, suggesting that the bacterium was circulating locally at some sites. This broad scale serological screening of whole seabird host communities provides critical information to assess disease emergence risks in the context of global change.

## Introduction

Tracking efficiently the health of threatened animal communities is needed to infer how global change may affect eco-epidemiological dynamics with consequences for biodiversity (Baker et al. 2022, Nova et al. 2022). Nevertheless, despite increasing recognition of the importance of infectious diseases in conservation biology (Smith et al. 2006, Young et al. 2017, Foufopoulos et al. 2022), investigations of disease outbreaks in wild populations often fail to determine what infectious agents were involved and fully assess their impacts (Childs et al. 2007, Ryser-Degiorgis 2013, Mazzamuto et al. 2021). Understanding the implications of infectious disease circulation in wild populations necessitates to explore what infectious agents with potential pathogenic effects are present or could emerge, whether they threaten populations, and at what spatial scales (Daszak et al. 2000). This requires sampling communities of potential host species that may share infectious agents, act as reservoirs (Plowright et al. 2019) or spatial spreaders for the disease (Gamble et al. 2020a, Clessin et al. 2025), this at both fine and broad spatial scales.

Disease monitoring in free-ranging wildlife populations is challenging and often relies on passive surveillance, consisting in the detection and characterization of abnormal mortality events (Ryser-Degiorgis 2013). This can lead to underreporting (e.g., in the case of the recent worldwide outbreaks of High Pathogenicity Avian Influenza (HPAI); Klaassen & Wille 2023). Alternatively, proactive targeted surveillance that relies on the systematic sampling of individuals independently of their apparent health state for detecting current infections (e.g., via PCR; Zamora et al. 2023) or evidence of past infections (e.g., via the detection of antibodies; Gilbert et al. 2013, Pepin et al. 2017) requires specific capture efforts and is often focused on a limited number of host species and sites (but see Wille et al. 2023). In order to explore what populations or segments of populations are exposed to potentially pathogenic agents, serological investigations are powerful, as they can inform on the history of exposure of individuals from the prevalence and level of antibodies against specific infectious agents in blood samples (Gilbert et al. 2013, Wilber et al. 2020). Various factors may affect antibody levels and their temporal persistence (Gilbert et al. 2013, Gamble et al. 2020b), but in many species, such as long-lived species faithful to specific breeding areas, antibody screening can be a particularly powerful tool to detect the occurrence and quantify the extent of the circulation of pathogens compared to attempts to directly detect the often-transitory infection of individuals (Lebarbenchon et al. 2015).

Seabirds that breed in dense colonies can be heavily impacted by epizooties caused by pathogens transmitted among socially interacting individuals (Vanstreels et al. 2023). This has been highlighted by recent outbreaks of HPAI, leading to massive local die-offs at colonies, but also widespread transmission among colonies, even at broad geographic scales (Boulinier 2023, Lane et al. 2024, Knief et al. 2024, Clessin et al. 2025). Because many seabirds aggregate seasonally on well-defined breeding areas, often on islands, they represent potentially useful systems to explore the processes involved in exposure to infectious agents at different spatial scales. Until the HPAI panzootic (Banyard et al. 2024, Clessin et al. 2025), few outbreaks among seabirds of the Southern Ocean have been the subject of specific investigations, apart for avian cholera(Uhart et al. 2018, Weimerskirch 2004).

Avian cholera is a widespread infectious disease caused by the bacterium *Pasteurella multocida* with significant economic impacts on poultry production (Singh et al., 2013). It also affects wild bird populations, notably anatids, for which massive die-offs have been observed in North America (Samuel et al., 2007; Thomas et al., 2007). Regarding seabirds (**Figure 1**), repeated outbreaks have been reported in common eiders (*Somateria mollissima*) on East Bay and Western Cape islands in Canada (Descamps et al., 2012; van Dijk et al., 2021; Waller & Underhill, 2007), as well as in various species of seabirds (skuas, giant petrels and penguins) over a long period of time on the Antarctic Peninsula (Parmelee, 1979; Leotta et al., 2003; 2006; Iervolino et al., 2025).

**Figure 1.**
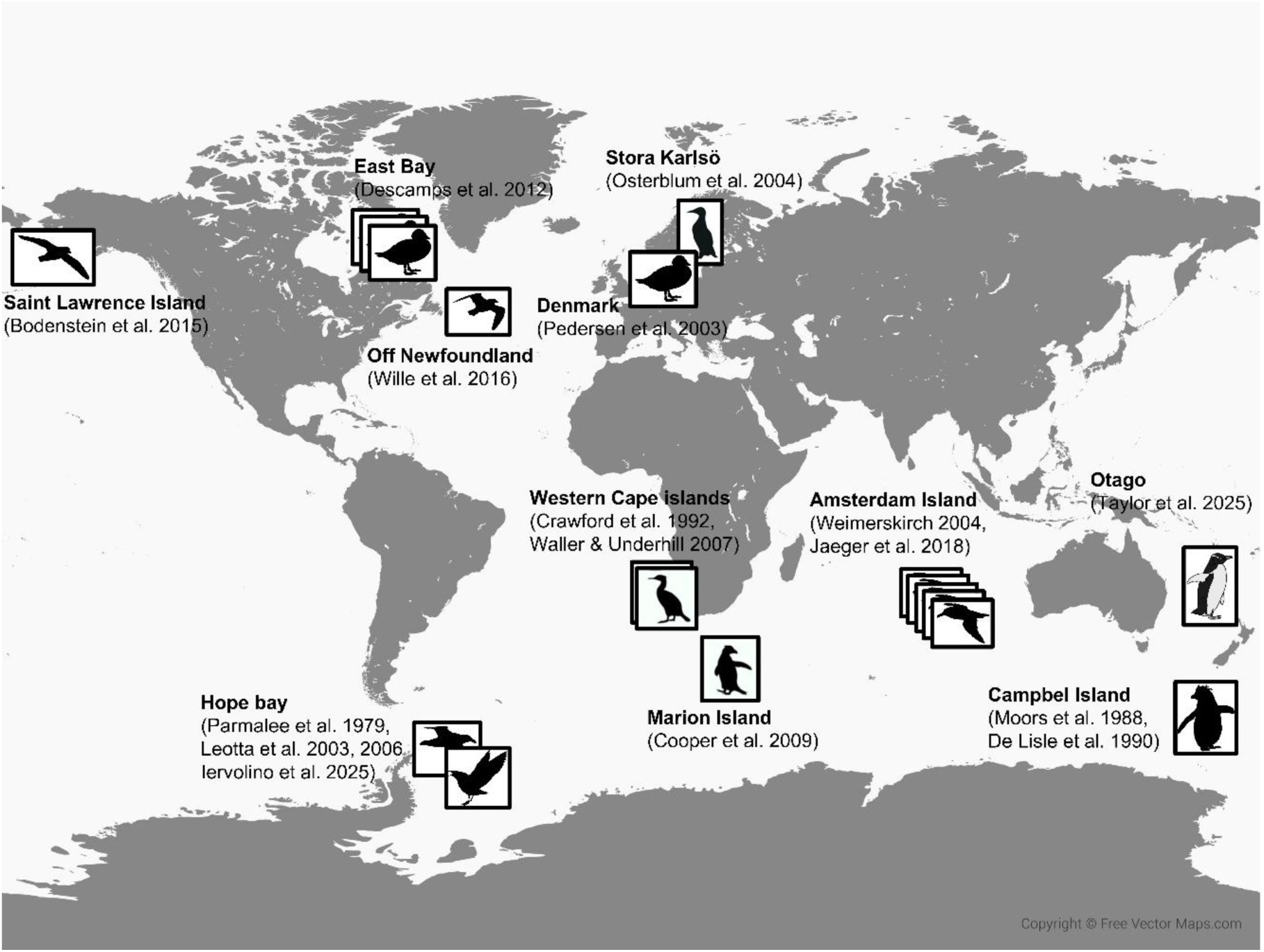
Map showing the main reported outbreaks of avian cholera in seabirds with references to published reports. Silhouettes of species outline the diversity of the affected hosts and references, while multiple silhouettes outline repeated reported outbreaks (Amsterdam Island, East Bay and Hope Bay).

A notable case is that of Amsterdam Island in the southern Indian Ocean, where this disease has caused high mortality rates among Indian yellow-nosed albatross (*Thalassarche carteri*) nestlings almost every year for several decades (Bourret et al., 2018; Jaeger et al., 2018; 2020). Sooty albatrosses (*Phoebetria fusca*) and Northern rockhopper penguins (*Eudyptes moseleyi*) have also been affected, and the disease threatens the small, endemic Amsterdam albatross population (*Diomedea amsterdamensis*) (Jaeger et al., 2018). The bacterium has repeatedly been isolated from the organs of dead seabirds on Amsterdam Island (Weimerskirch, 2004; Jaeger et al., 2018, 2020), enabling the whole genome of the main circulating strain to be sequenced (Hashish et al., 2023). This strain has been identified as ST91 using the multilocus sequence typing (MLST) method (Jolley et al., 2018). ST91 was recently identified as being responsible for a massive die-off of Adelie penguins on the Antarctic Peninsula (Iervolino et al., 2025). Another strain, ST110, was also detected in a small number of necropsied albatrosses on Amsterdam Island (Tornos et al., unpublished). Both STs are Capsular A, Serotype 1, ssp. *P. multocida gallicida*.

The pathogenicity of *P. multocida* typically varies among and within host species (Blanchong et al., 2006). For example, avian cholera primarily affects albatross juveniles on Amsterdam Island (Bourret et al., 2018), while it has been reported to impact the survival of adult common eider (Descamps et al. 2012) and gulls in North America (Wille et al. 2016) as well as brown skuas (*Stercorarius antarcticus*), Southern giant petrels (*Macronectes giganteus*) and Adelie penguins (*Pygoscelis papua)* in Antarctica (Parmalee et al. 1979, Leotta et al. 2003, 2006). Conversely, some populations or species sustain lower mortality rates and act as chronic carriers of *P. multocida*, including certain North American geese (Samuel et al., 2005) and brown skuas on Amsterdam Island (Gamble et al., 2020a; Lamb et al., 2023). The diversity of strains may also be responsible for the observed differences in pathogenicity between host populations (Christensen et al., 1998; Harper et al., 2006).

Exposure of birds to *P. multocida* can lead to acute or chronic infections. In acute infections, few clinical signs are observed before death and histopathological signs are dominated by general septicaemic lesions (Christensen & Birgaard 2000). In most cases, transmission is thought to be either by inhalation of a bacteria-contaminated aerosol or through the consumption of contaminated water or food (Botzler 1999). Many *Pasteurella* species are opportunistic pathogens that can cause endemic disease and are associated increasingly with epizootic outbreaks (Wilson & Ho 2013). Zoonotic transmission to humans usually occurs through animal bites or contact with nasal secretions, with *P. multocida* being the most prevalent isolate observed in human infections. As often, it is challenging to disentangle individual exposure from susceptibility (Sweeny & Albery 2023). In the case of Indian yellow-nosed albatrosses on Amsterdam Island for instance, nestlings are most severely affected, but it is unclear whether this is due to higher susceptibility or greater exposure compared to adults (Gamble et al. 2019, Jaeger et al. 2020). Moreover, in some populations hosting such a bacterium commensally, changes in environmental conditions have been identified as the cause of massive die-offs (Koch et al. 2018). Determining whether widely dispersed seabird populations are exposed to *P. multocida* by detecting antibodies is thus an important step to infer whether the disease could emerge as a broad scale threat.

Many seabird species are threatened globally, especially groups like petrels, albatrosses and penguins (Dias et al. 2019), and infectious diseases represent a neglected threat with a potential to increase dramatically with environmental changes. In this context, seabird species with apex scavenging or predatory habits could be used as sentinels (Halliday et al. 2007) as they may be at greater risk of exposure to infectious agents while feeding on infected individuals or carrion. Within the relatively simple social networks made of seabird colonies, sentinels could be used to efficiently assess where pathogens are present and whether they may emerge as substantial threats to populations (Bai et al. 2017). In return, such species may also contribute to the spread of infectious agents (Gamble et al. 2020a). More broadly, seabird communities are valuable systems for the study of infectious disease dynamics. Although individuals in seabird colonies can number over hundreds of thousands, the numbers of species are usually relatively small (at most tens of species), with well-defined distributions in space (Nelson 1980), which simplifies field sampling. Populations of similar species can be sampled at multiple sites, which could then represent spatial replicates, allowing reporting more robust results than if samples were obtained from a limited number of locations.

By conducting broad-scale antibody screening of seabird communities in the southern Indian and Atlantic oceans to quantify exposure to *P. multocida* (**Figure 2**), we assessed the spatial scale at which such an infectious agent may be a threat, and tested whether scavenging or avian predator species, notably brown skuas, giant petrels and lesser sheathbills (*Chionis minor*), could be used as sentinels to track its circulation. We used complementary approaches to confirm exposure to the bacterium and we predicted that petrels and skuas could inform us of exposure at broad scales, while the non-migratory lesser sheathbill (Verheyden & Jouventin 1991) could provide more local information. The results stress that considering fine-to-broad spatial scales and hosts with contrasted ecologies is critical when aiming at improving the monitoring of emergence risk of multi-host diseases in wild animal communities.

**Figure 2.**
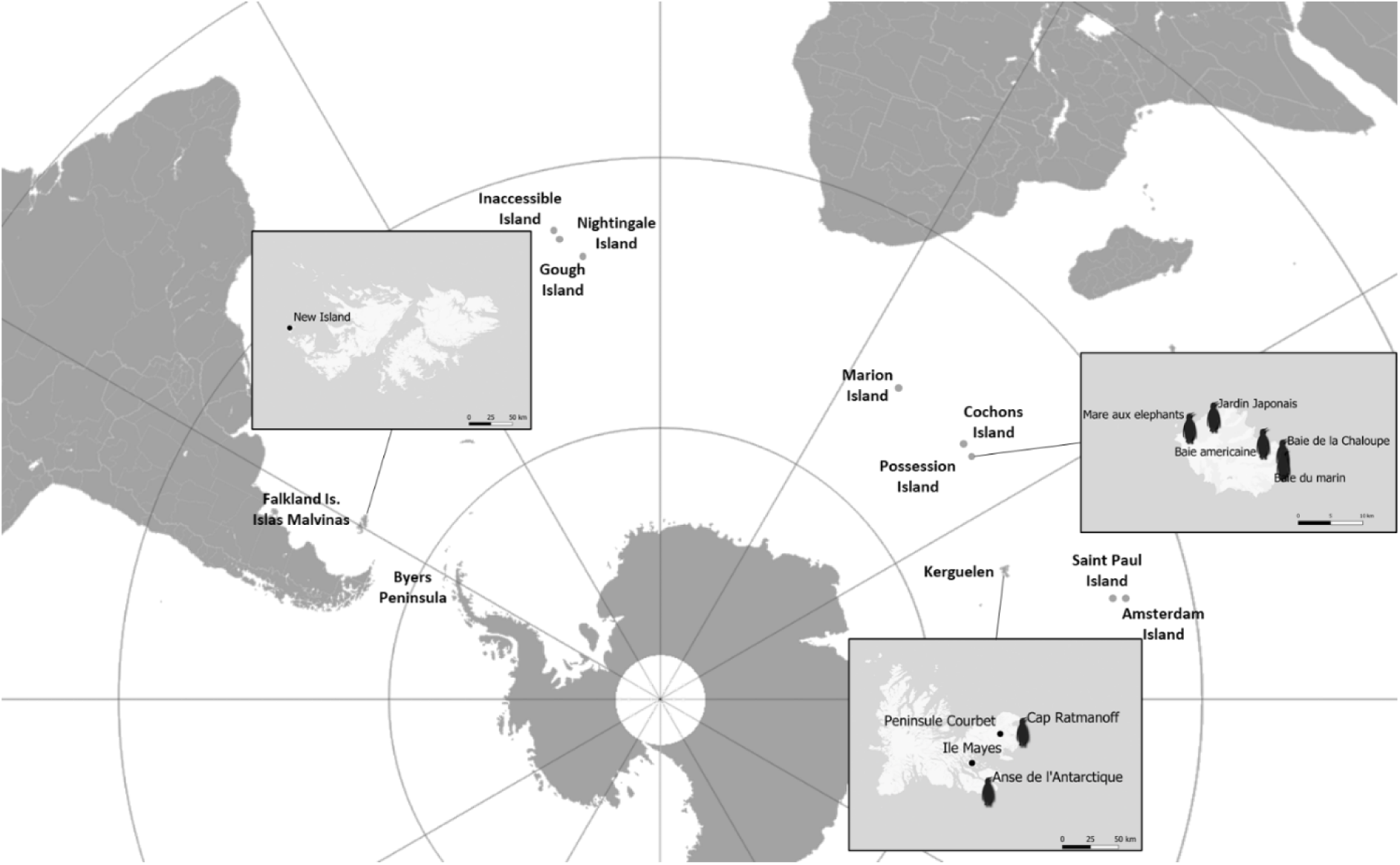
Sampling locations. Inserts: details of the sampling locations within Crozet and Kerguelen archipelagos, where series of sites were sampled. The main sampling sites, which hosts large to very large king penguin colonies, are marked with king penguin symbols and their names are given. The small dots show the location of New Island, in the Falkland Islands/Islas Malvinas, and of Mayes Island and Péninsule Courbet in Kerguelen archipelago.

## Results

### Detection of broad scale exposure to *Pasteurella multocida*

Among the 2634 blood samples analyzed, antibodies against *P. multocida* were detected in 14 of the 18 seabird species. High exposures of several seabird species to *P. multocida* were found (high microagglutination (MAT) titres and high seroprevalences; **Table 1**), but they differed greatly among host species and geographic areas (**Figure 3**; see details of modelling results per species in **Supplement S1**). In particular, widely roaming scavengers and avian predators like brown skuas and giant petrels showed very high titres (**Figure 3, 4, 5**) and seroprevalences (> 0.9; **Figure 3, 6**). When grouped according to their ecologies, apex predators and scavengers showed, as predicted, higher seroprevalences and mean titres than the other groups (**Table 2**). Most Northern giant petrels (*M. halli*) showed very high titres and all were seropositive (**Figure 3, 5**). Southern giant petrels showed also high titres and prevalence, with slightly lower values for the Antarctic Peninsula (**Figure 3, 5**). Brown skuas, a species that breeds on all austral sites sampled, showed variations among sites, with the highest mean titres found on Amsterdam Island (where recurrent outbreaks have been causing mass nestling mortality events), followed by Gough, Marion, Crozet and Kerguelen Islands; conversely, very low prevalence and a mean titre lower than 10 was found for brown skuas on New Island, Falklands Islands (**Figure 4, 6**). No sex effect was detected in any species (**Supplement S1**).

**Figure 3.**
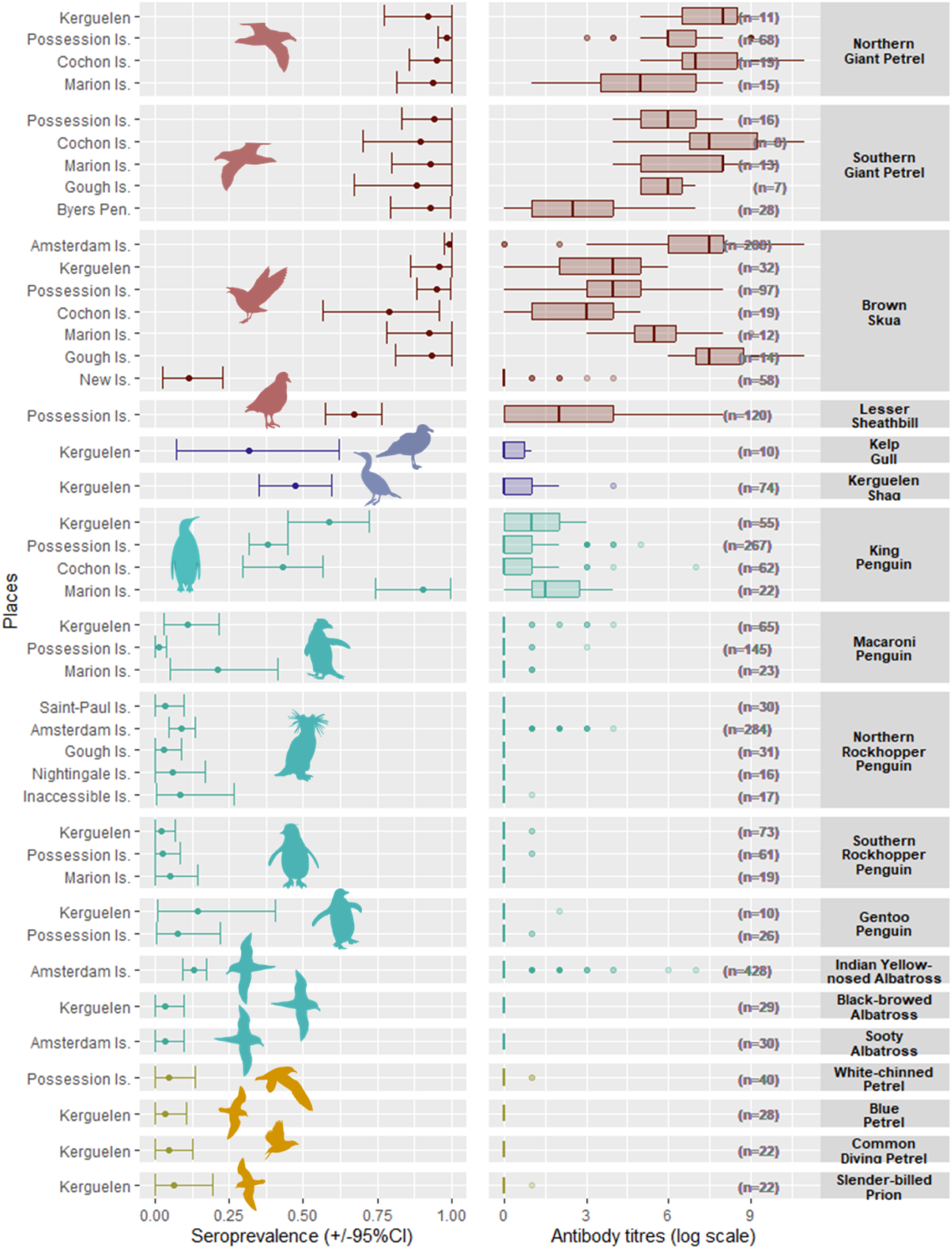
Seroprevalences and mean antibody titres against *Pasteurella multocida* in the sampled species and sites. Titres above 1/10 (i.e. 0 on the log scale) were considered seropositive. Species are grouped as (i) in red, the apex predators and scavengers, (ii) in blue, the ground nesting inshore meso predators, (iii) in green, the ground nesting offshore meso predators and (iv) in yellow, the burrow nesting offshore meso predators (Supplement S2). The scientific names of the species are given in legend of **Table 1**. Bars display 95 % confidence intervals.

**Figure 4:**
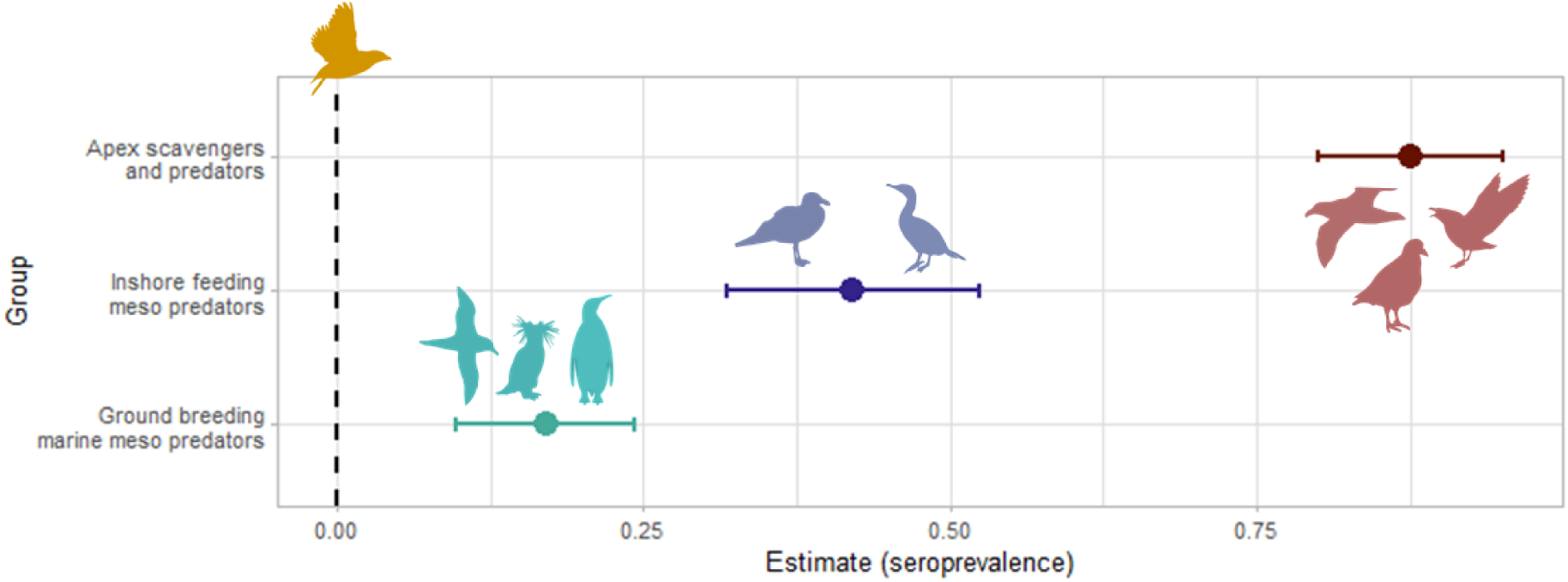
Standardised estimates of the parameters of the generalised linear models per group in relation to the seroprevalence levels. The colours are representing the respective groups, (i) in red, the apex predators and scavengers, (ii) in blue, the ground nesting inshore meso predators, (iii) in green, the ground nesting offshore meso predators, (iv) in yellow, the burrow nesting offshore predators, taken as the reference value. The black dashed line highlights the position of 0 on the x-axis. The variable is significant if the confidence interval excludes 0. More details in **Table 2**.

**Figure 5.**
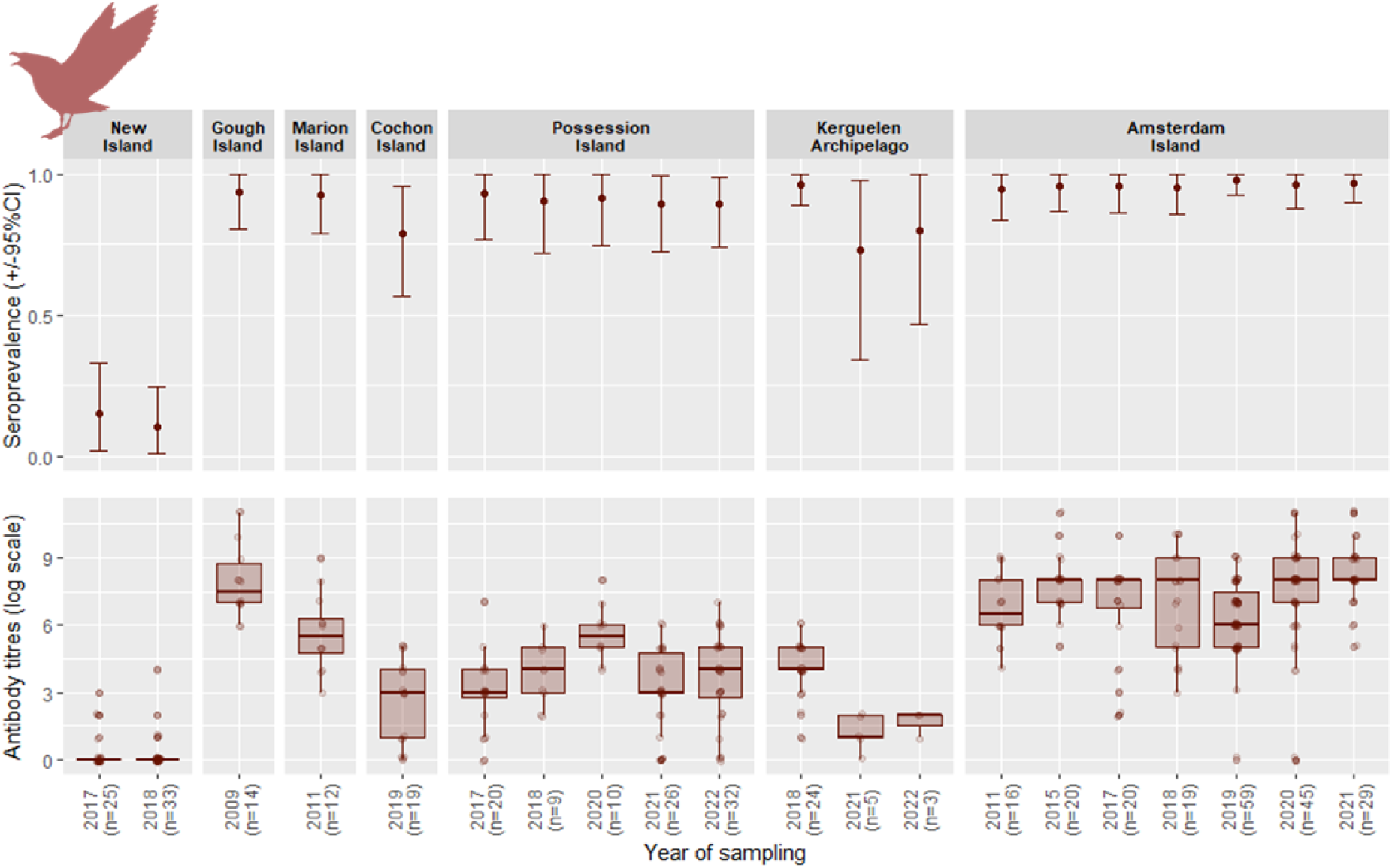
Seroprevalence and mean antibody titres (bottom) against *Pasteurella multocida* in brown skuas sampled on islands of the southern Atlantic and Indian oceans over several years (sample size in brackets for each year and site). Titres above 1/10 (i.e. 0 on the log scale) were considered seropositive. High antibody titres were observed at all sites but New Island, suggesting a broad spatial exposure of populations of this species, but not on the southern tip of South America. On Amsterdam Island, recurrent outbreaks of avian cholera have been responsible for massive die-offs of Indian yellow-nosed albatross chicks, on which the local skuas scavenge, which likely explains their very high seroprevalence and antibody titres. It is unclear where the skuas from Marion Island, Gough Island and Crozet and Kerguelen Islands get exposed to *Pasteurella multocida*, although the variable titres and seroprevalences detected in king penguins in all large colonies where they could be sampled (n = 7 locations, on Marion Island, Possession Island, Île aux Cochon and Kerguelen archipelago) could suggest that brown skuas may be exposed when scavenging on such a species.

**Figure 6.**
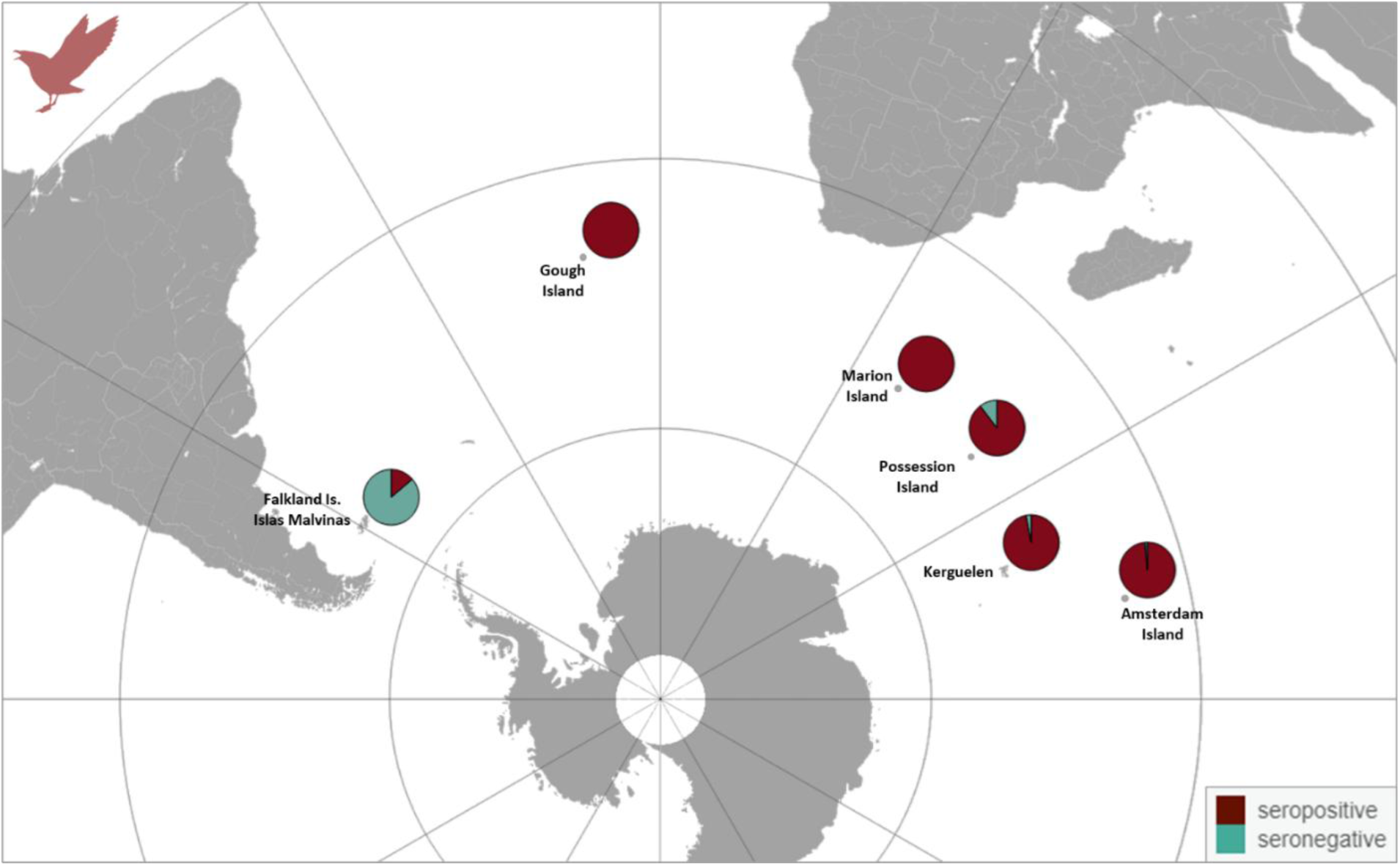
Map showing the seroprevalence of brown skuas against *Pasteurella multocida* for all sampled locations for all years. The pie charts represent in red the proportion of seropositive individuals. Those results highlight the broad spatial exposure of this scavenger species and the importance of surveys at large spatial scales.

**Table 1.**
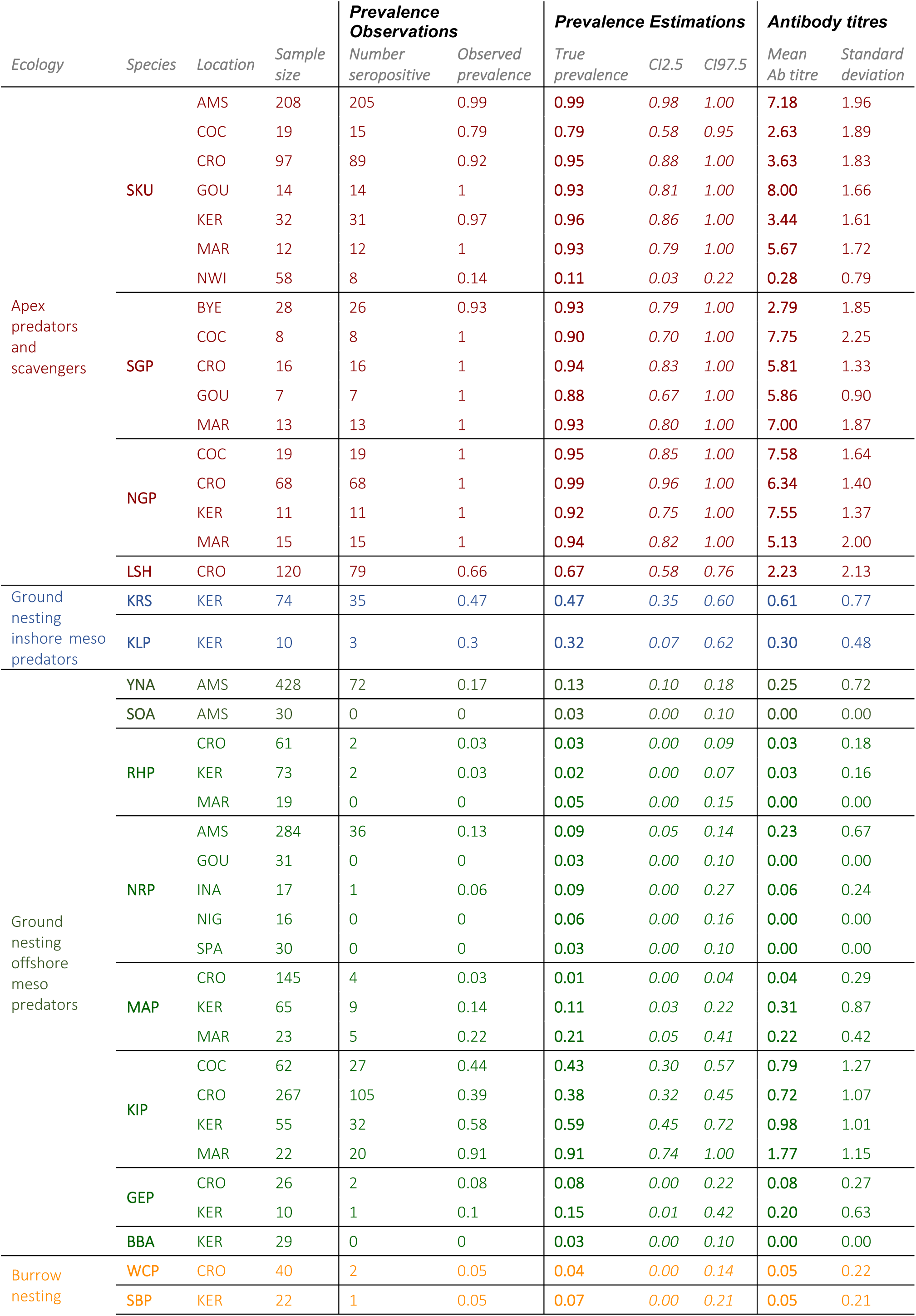
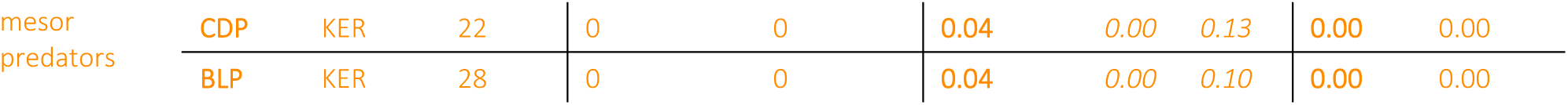
Seroprevalences observed and calculated (true prevalence) with 95% confident interval and mean antibody titres with standard deviation as a function of species and locations of sampling (threshold titre for seropositivity <= 1/10 or > 0 on the log scale). Species are grouped as (i) the apex predators and scavengers, (ii) the inshore meso predators, (iii) the ground nesting offshore meso predators and (iv) the burrow nesting offshore meso predators (**Supplement S2**). Each group is represented by a color, respectively (i) red, (ii) blue, (iii) green and (iv) yellow. Species are: Northern giant petrel *Macronectes halli* (NGP), Southern giant petrel *Macronectes giganteus* (SGP), brown skua *Stercorarius antarcticus* (SKU), lesser sheathbill *Chionis minor* (LSH), kelp gull *Larus dominicanus* (KLP), king penguin *Aptenodytes patagonicus* (KIP), Kerguelen shag *Phalacrocorax verrucosus* (KRS), macaroni penguin *Eudyptes chrysolophus* (MAP), Northern rockhopper penguin *Eudyptes moseleyi* (NRP), Southern rockhopper penguin *E. chrysocome* (RHP), gentoo penguin *Pygoscelis papua* (GEP), Indian yellow-nosed albatross *Thalassarche carteri* (YNA), black-browed albatross *T. melanophris* (BBA), sooty albatross *Phoebetria fusca* (SOA), white-chinned petrel *Procellaria aequinoctialis* (WCP), blue petrel *Halobaena caerulea* (BLP), common diving petrel *Pelecanoides urinatrix* (CDP), slender-billed prion *Pachyptila belcheri* (SBP). Codes for the location names are: AMS for Amsterdam Island, COC for Île aux Cochons, CRO for Île de la Possession in Crozet archipelago, GOU for Gough Island, KER for Kerguelen archipelago, MAR for Marion Island in Prince Edwards Islands archipelago, NYI for New Island in the Falkland Islands, INA for Inaccessible Island, NIG for Nightingale Island and SPA for Saint Paul Island.

**Table 2.**
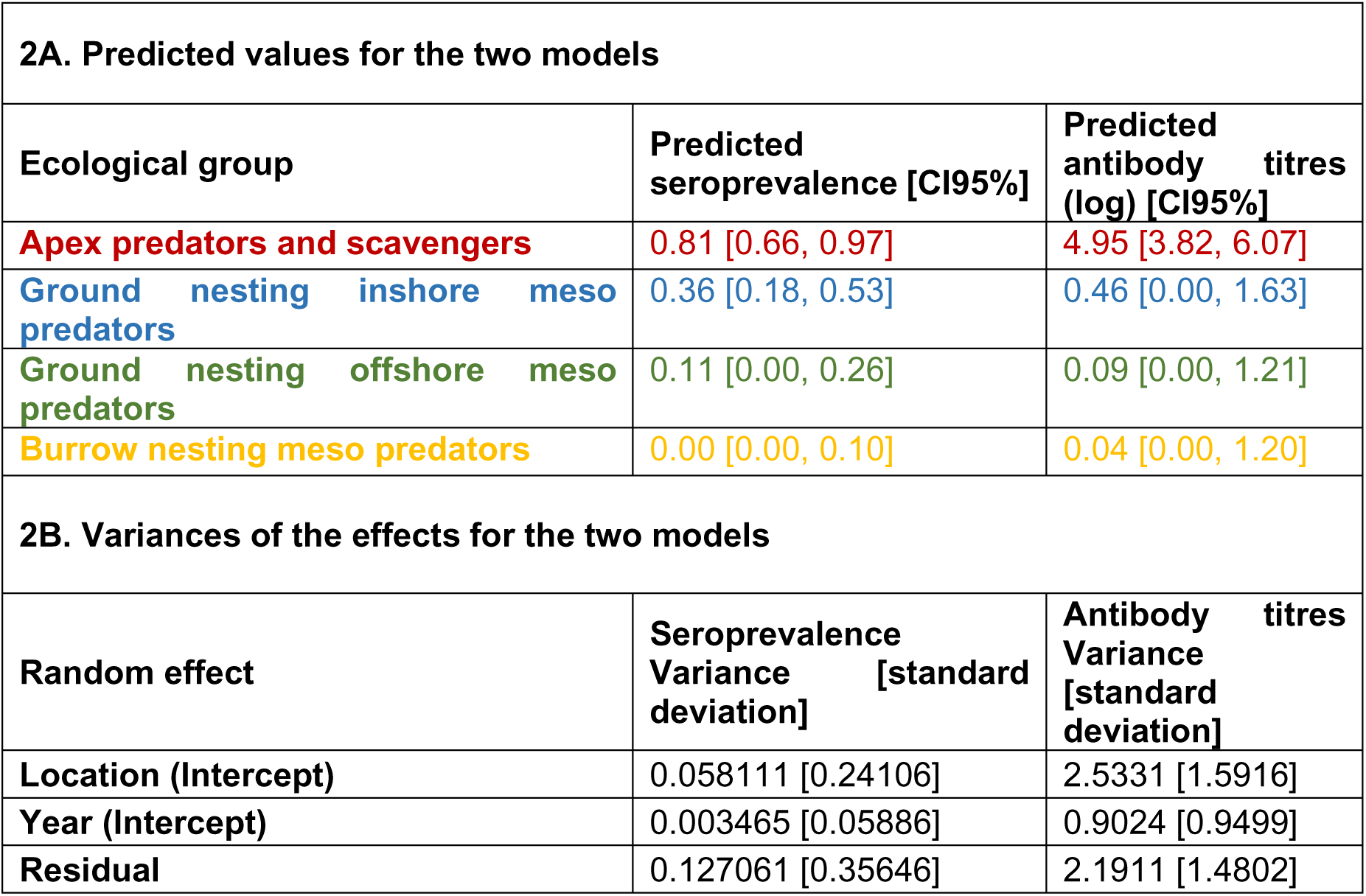
Output of the GLMM modelling of the seroprevalence and antibody titres as a function of the ecology groups, accounting for random location and year. 95% confidence interval for each ecological group are given in brackets. Sub-table 2A presents the predicted seroprevalences and antibody titres for each group for the two models, and sub-table 2B presents the variances of the effects for the two models.

| 2A. Predicted values for the two models |  |  |
| --- | --- | --- |
| Ecological group | Predicted seroprevalence [CI95%] | Predicted antibody titres (log) [CI95%] |
| Apex predators and scavengers | 0.81 [0.66, 0.97] | 4.95 [3.82, 6.07] |
| Ground nesting inshore meso predators | 0.36 [0.18, 0.53] | 0.46 [0.00, 1.63] |
| Ground nesting offshore meso predators | 0.11 [0.00, 0.26] | 0.09 [0.00, 1.21] |
| Burrow nesting meso predators | 0.00 [0.00, 0.10] | 0.04 [0.00, 1.20] |
| 2B. Variances of the effects for the two models |  |  |
| Random effect | Seroprevalence Variance [standard deviation] | Antibody titres Variance [standard deviation] |
| Location (Intercept) | 0.058111 [0.24106] | 2.5331 [1.5916] |
| Year (Intercept) | 0.003465 [0.05886] | 0.9024 [0.9499] |
| Residual | 0.127061 [0.35646] | 2.1911 [1.4802] |

### The particular case of Amsterdam Island

On Amsterdam Island, where recurrent outbreaks of avian cholera have been killing many Indian yellow-nosed albatross nestlings and where the highest seroprevalences and titres were found for brown skuas, relatively low levels were detected in some adult Indian yellow-nosed albatrosses and the overall seroprevalence among adults was low (< 20%; **Table 1**, **Figure 3**). Northern rockhopper penguins also showed very low seroprevalence (while no evidence of antibodies against *P. multocida* was found in individuals of that species sampled on Saint Paul Island and Gough, Inaccessible and Nightingale Islands, Tristan da Cunha; **Figure 3**). No antibody was found in a small sample of adults of the population of sooty albatrosses from Amsterdam Island (**Figure 3**, **Table 1**).

### Variability in exposure across seabird communities

At sampled sites, 38-91% of king penguins (*Aptenodytes patagonicus*), lesser sheathbills and Kerguelen shags (*Phalacrocorax verrucosus*) showed evidence of antibodies against *P. multocida* (**Table 1**, **Figure 3, 7, 8**). For king penguins, a higher mean titre was found for Marion Island compared to the other sites. Within Possession Island, Crozet, the only location where king penguins could be sampled several years, a year effect was detected in the mean titre. For lesser sheathbills, annual residents on the island, mean titres differed among colonies and years within Possession Island. Their seropositivity at the main king penguin colonies suggests that, as a year-round resident species, they could be a good sentinel at that scale. For colonies where both king penguins and lesser sheathbills were sampled on Possession Island, strong spatial and temporal variation was detected for both species, but the temporal variations were not correlated (**Figure 7**; r = 0.26, P = 0.54), and mean titres were higher for lesser sheathbills than for king penguins (**Figure 8 & Figure 9**). Conversely, Southern rockhopper penguins *Eudyptes chrysocome* (ssp*. filholi* and *chrysocome*), macaroni (*E. chrysolophus*) and gentoo (*Pygoscelis papua*) penguins showed only a few seropositive individuals (**Figure 3**). Less than 5% of individuals were found positive in the four burrowing petrel species (white-chinned petrel (*Procellaria aequinoctialis)* sampled on Possession Island, Crozet, and blue petrel (*Halobaena caerulea)*, common diving petrel (*Pelecanoides urinatrix)* and slender-billed prion (*Pachyptila belcheri)* on Mayes Island, Kerguelen; **Table 1**).

**Figure 7.**
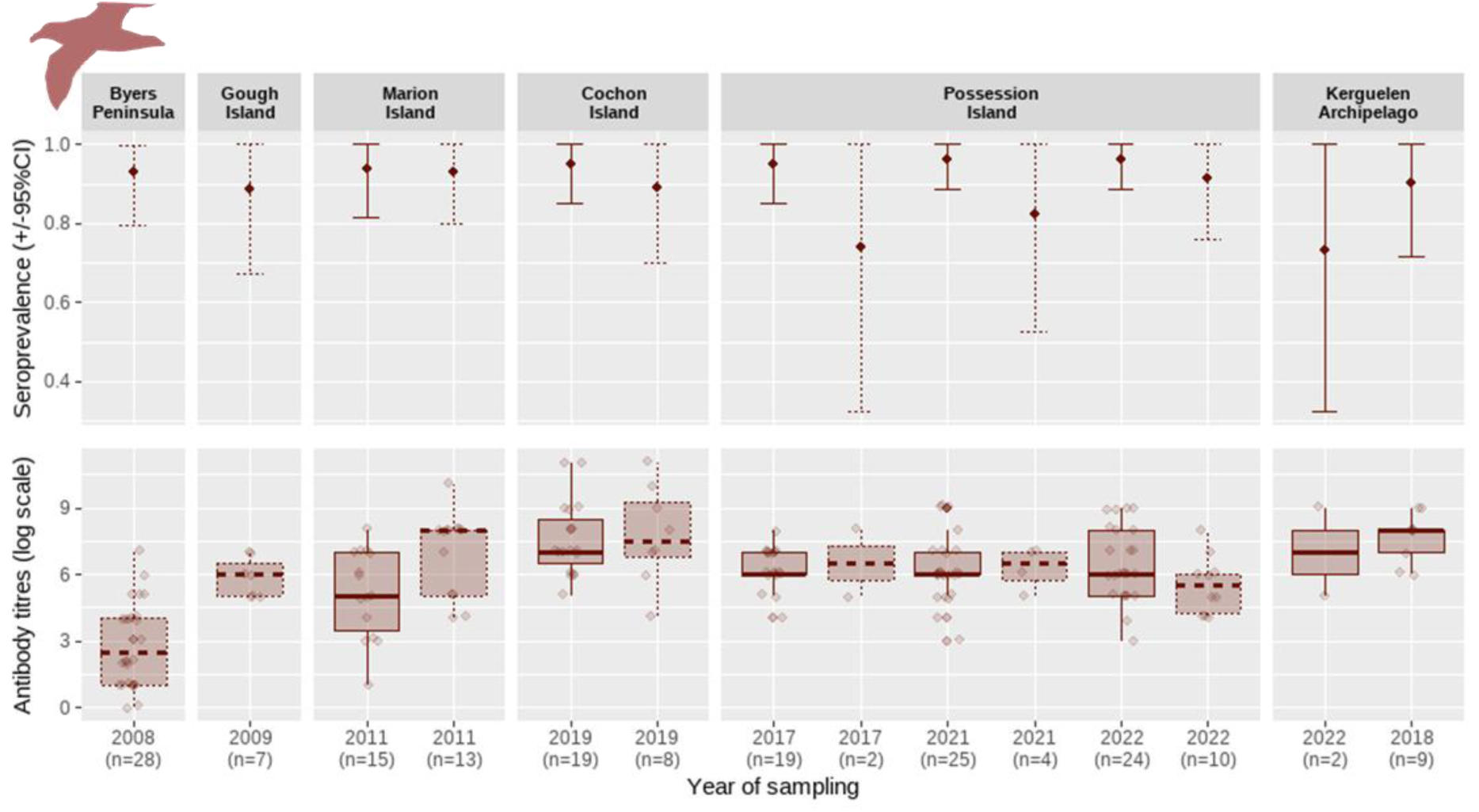
Seroprevalence and mean antibody titres against *Pasteurella multocida* of the Northern and Southern giant petrels (respectively plain and dashed lines) at all sampled locations, from the Antarctic peninsula to Kerguelen archipelago. Both species of scavengers show very higher titres and seroprevalences.

**Figure 8.**
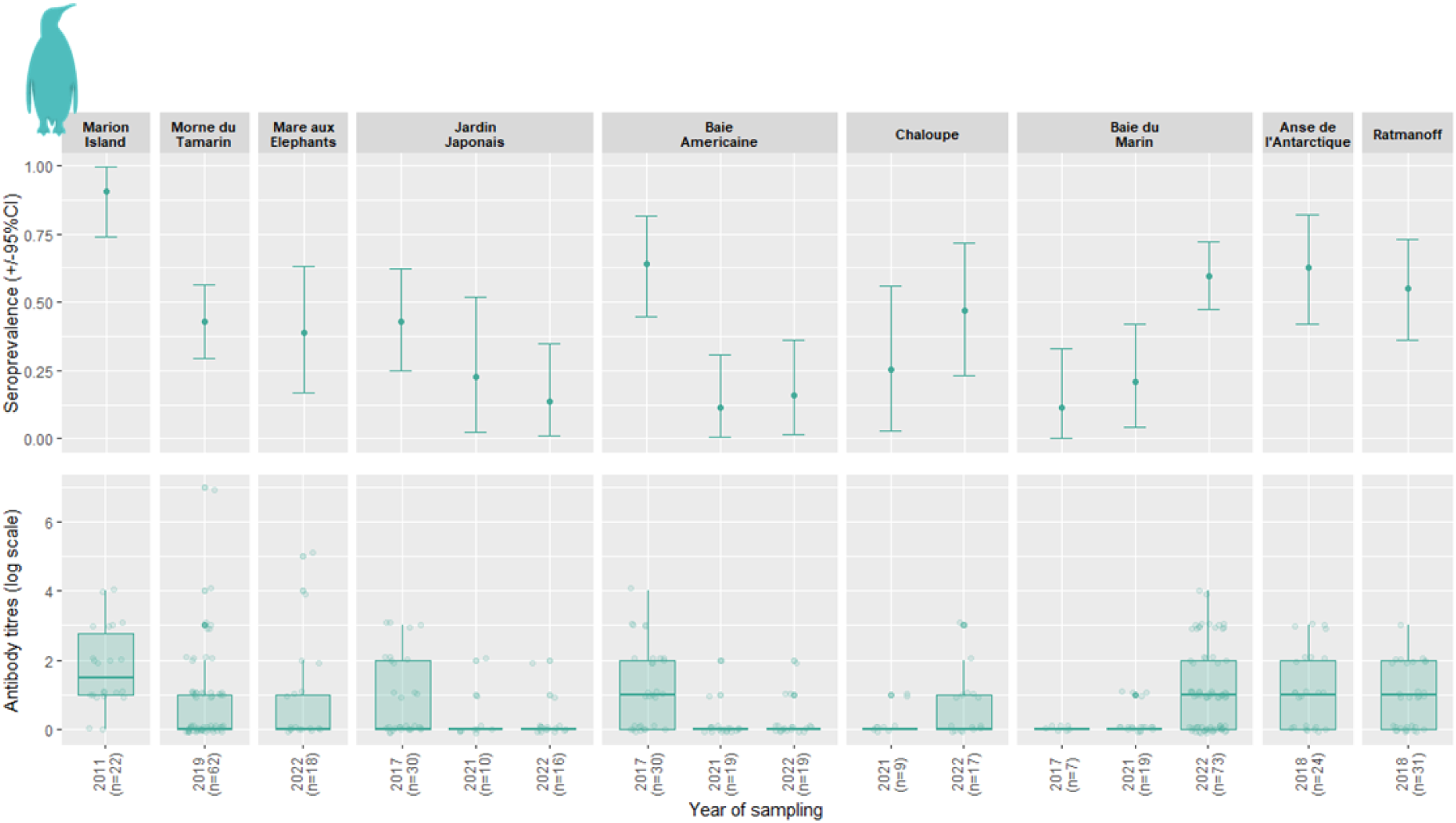
Seroprevalence and mean antibody titres against *Pasteurella multocida* in king penguin colonies sampled in the southern Indian Ocean. Variable proportions of king penguins have antibodies against *Pasteurella multocida* and data for series of years show that there is a relative temporal stability of the results among years.

**Figure 9.**
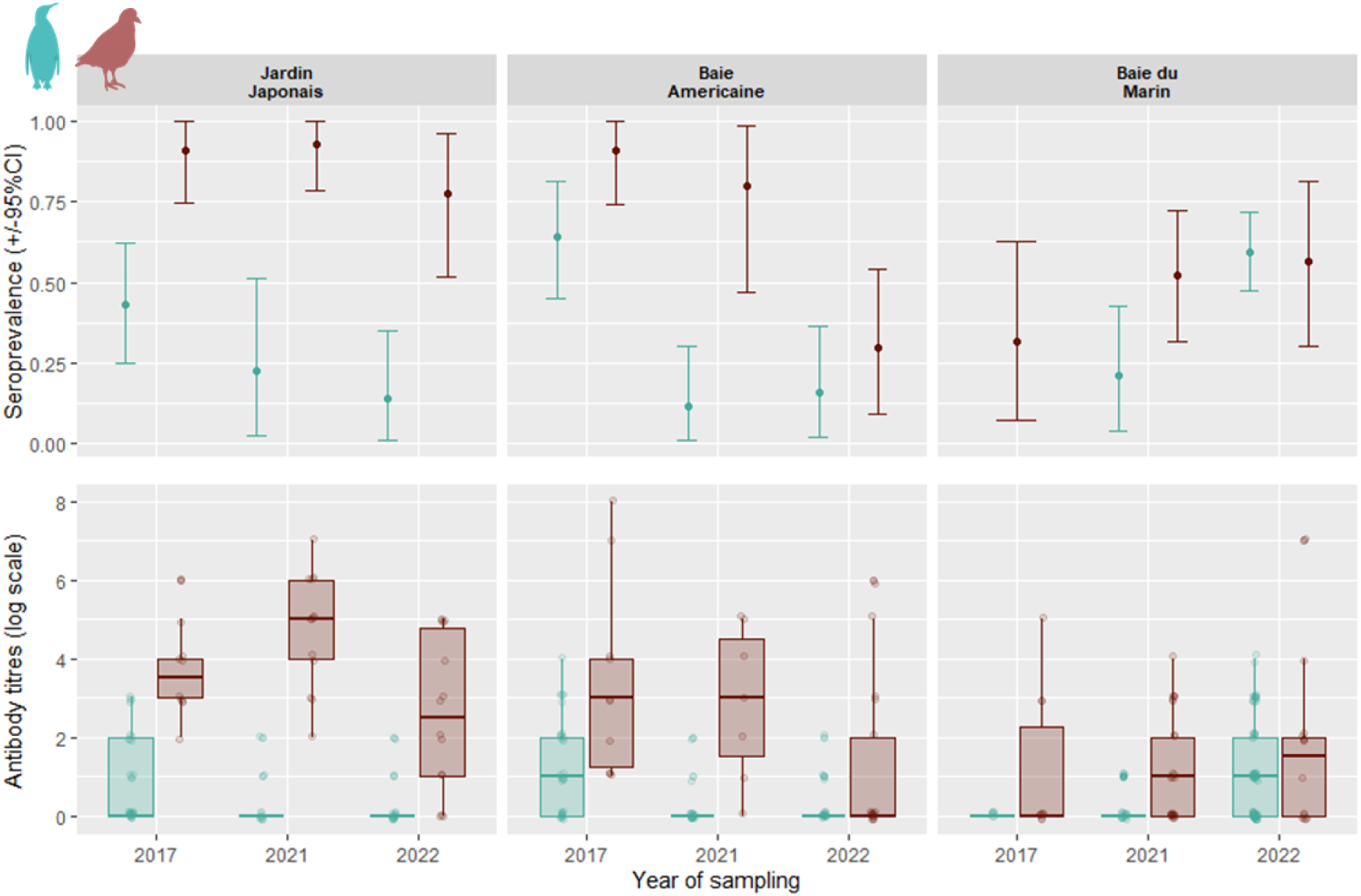
Seroprevalence (estimated prevalence) and mean antibody titres against *Pasteurella multocida* in king penguin and lesser sheathbill within Possession Island, Crozet archipelago, where 3 colony sites were sampled for both species.

Overall, the patterns were strongly consistent among islands between groups of the same species (**Figure 3**, **Table 1**). Island-scale patterns were found highly repeatable between years where they could be investigated, although at finer scale, between-colony differences could be detected, e.g., among king penguins on Crozet Islands (**Figure 7**, **Table 3**).

**Table 3:** Summary statistics of the multinomial logistic regression and analysis of variance of the effects of host species, location and years on, respectively, the seroprevalence and mean antibody titres against *Pasteurella multocida* in lesser sheathbills (LSH) and king penguins (KIP). Colonies at 3 sites were sampled for both species on Île de la Possession Island, Crozet archipelago: Jardin japonais (JJAP); Baie Américaine (BUS); Baie du Marin (BDM). Significant p-values (<0.01) are written in bold italic. Abbreviation: CI, confidence interval. Seroprevalence and mean antibody titres were higher in lesser sheathbills than in king penguins. The global seroprevalence was lower for the year 2021 and the mean antibody titres are higher in JJAP (see text).

| ***Predictors*** | ***Estimates*** | ***2.5% lower CI*** | ***97.5% upper CI*** | ***p-value*** |
| --- | --- | --- | --- | --- |
| ***<u>Seroprevalence</u>*** |  |  |  |  |
| Intercept | 0.47 | 0.34 | 0.59 | **0.0001** |
| Species: Lesser Sheathbill | 0.28 | 0.17 | 0.39 | **0.0001** |
| Year: 2021 | -0.19 | -0.33 | -0.05 | **0.0084** |
| Year: 2022 | -0.09 | -0.22 | 0.03 | 0.1387 |
| Colony: Jardin Japonais | 0.02 | -0.11 | 0.15 | 0.7673 |
| Colony: Baie Americaine | -0.07 | -0.19 | 0.06 | 0.3181 |
| ***<u>Mean antibody titres</u>*** |  |  |  |  |
| Intercept | 0.57 | 0.18 | 0.97 | **0.0047** |
| Species: Lesser Sheathbill | 1.49 | 1.15 | 1.83 | **0.0001** |
| Year: 2021 | -0.37 | -0.81 | 0.08 | 0.1064 |
| Year: 2022 | -0.25 | -0.65 | 0.15 | 0.2258 |
| Colony: Jardin Japonais | 0.68 | 0.27 | 1.08 | **0.0010** |
| Colony: Baie Americaine | 0.14 | -0.25 | 0.53 | 0.4844 |

### Cases of avian cholera mortalities in king penguins on Crozet Islands

No large-scale abnormal die-offs of seabirds had been recorded on Crozet and Kerguelen Islands until the emergence of HPAI on Possession Island, Crozet, in October 2024 (Clessin et al., 2025). However, isolated mortalities have prompted necropsies and sample collection from recently dead individuals as part of a long-term disease monitoring program. One such necropsy was conducted on an adult king penguin found dead in April 2022 at the Baie du Marin colony, near the research station on the Possession Island, Crozet. Histopathological investigation revealed lesions consistent with a *Pasteurella multocida* infection. Corresponding samples from several organs tested positive for PCR, strongly suggesting that avian cholera was the cause of death (see **Supplement S4** for details). In November 2023, another freshly dead adult king penguin was necropsied at the same colony, and bacterial isolates were obtained from swabs of all internal organs using a selective culture medium. PCR testing of an isolate again suggested that avian cholera was the cause of death, although no histological data are available (**Supplement S4**).

## Discussion

### Widespread but heterogeneous exposure of seabird communities

High seroprevalences and antibody titres against *Pasteurella multocida* were found at all locations but the Falkland Islands in three wide roaming species that could be considered as efficient sentinels at a broad spatial scale, the Northern and Southern giant petrels and brown skua. Our serology approach does not provide information on whether the bacterium is pathogenic at various locations, but it does suggest that individuals of various species get infected and mount immune responses against *Pasteurella multocida*. Our report of two suspected mortalities of adult king penguins due to avian cholera on Crozet archipelago suggests that the bacterium could be responsible for low-intensity die-offs outside the large-scale die-offs reported in the literature (Jaeger et al. 2020, Iervolino et al. 2025). Alerting on the presence of the bacterium can be important to focus specific research efforts, but also to stress the importance of biosecurity issues for people handling seabirds or visiting colonies. At some locations, variable proportions of king penguins, lesser sheathbills and shags showed evidence of antibodies against *P. multocida*, suggesting local circulation to the bacterium. It is not informing us on whether the bacterium has been responsible for some die-offs of individuals, but it stresses the need for more surveillance, notably via conducting necropsies on dead individuals that can be found in the field. Several species, notably burrow-nesting petrels, showed almost no sign of exposure to *P. multocida*. On Amsterdam Island, where regular outbreaks of avian cholera have been documented, notably by necropsies followed by *P. multocida* cultures and positive PCR from internal organs (Weimerskirch 2004, Jaeger et al. 2020), low but non-negligible seroprevalences and antibody titres were found in the locally abundant and highly affected Indian yellow-nosed albatross (Gamble et al. 2019), while seroprevalence was almost 100% each year in adult brown skuas (Gamble et al. 2020a, Lamb et al. 2023). Northern rockhopper penguins showed low seroprevalence, despite possibly suffering from outbreaks of the disease on Amsterdam Island (Jaeger et al. 2018). Overall, the patterns were repeatable between years at locations where they were investigated; the same set of species showed high seroprevalences and antibody titres. At fine scales, some temporal and spatial variability was detected within Crozet Islands for king penguins and lesser sheathbills, indicating underlying local dynamics. At larger scales, higher seroprevalences and antibody titres were detected in brown skuas on Amsterdam Island, while they were lower at Kerguelen, Crozet and Marion Islands. In the Falkland Islands, almost no seropositive skua individuals was detected over a number of years. For Southern giant petrels, seroprevalence was close to 1 at all sites, but a lower mean antibody titre was detected at Byers, on the Antarctic Peninsula, suggesting less frequent exposure of the individuals sampled at this site. Infections by the bacterium *P. multocida* are known to be responsible for recurrent and large outbreaks of avian cholera in seabirds on subtropical Amsterdam Island, and sporadic outbreaks of the disease have been reported at other locations of the Southern Hemisphere, in South Africa, New Zealand and Antarctica (**Figure 1**). In particular, recent avian cholera outbreaks have been reported affecting penguins (Taylor et al. 2025, Iervolino et al. 2025). In this context, our serological results stress that it would be important to explore via more systematic necropsy investigations whether avian cholera might be responsible for some unreported morbidities or mortalities at some sites.

The results also raise a series of questions, which we discuss below: Are the strains circulating in seabird populations outside of Amsterdam Island and the Antarctic Peninsula pathogenic and related? Do mortality effects and immune responses differ among species? Is the bacterium maintained at some sites by introduced mammal species and what environmental factors play a role in the eco-epidemiological dynamics? It would also be important to know how the dynamics of the level of antibodies against *P. multocida* may vary among seabird species and how this could affect inferences from serological surveys.

### Scale of movements and sentinel values of scavenger and predatory species

Depending on the movement habits of the seabird host species and their life histories, it may be possible to infer where seropositive individuals were infected by the bacterium and thus the scale at which some species may be used as sentinels (Boulinier et al. 2016). In particular, giant petrels and brown skuas, which are migrants visiting various locations throughout their annual cycle (Thiers et al. 2014, De Souza Petersen et al. 2017, Delord et al. 2018), may be exposed to the bacterium at sites far from their breeding locality. It is unlikely that infected seabirds from Amsterdam Island represent a significant source of exposure of giant petrels as they are only scarce visitors to the island, which suggests other potential sources. The patterns of seropositivity of lesser sheathbill and king penguins, both confined to the sub-Antarctic and seropositive at location away from Amsterdam Island, suggest local and likely repeated exposure at some sites, notably on Possession Island, Crozet, because the expected turnover of individuals may not explain persistent seroprevalences at a location without novel exposure. It is unclear though whether this may be due to an enzootic circulation or repeated introductions of strains or rare spreading events. Super-spreading events (Lloyd-Smith et al. 2005) could be particularly important given the highly social habits of many species, the strong spatial structure of the system and the opportunities for movements by some individuals, such as skuas and giant petrels, among locations. Anyway, the results stress that a scavenging and yearly resident species like the lesser sheathbill (Verheyden & Jouventin 1991) may represent an especially interesting sentinel at the scale of an archipelago like Crozet.

### Integrating serology with complementary data to infer transmission within and among host populations

Inferring connectivity among host populations via data on movements of individuals can be useful, but evidence of visits of colonies does not imply transmission events. While roaming of skuas or giant petrels at the scale of Crozet archipelago could contribute to epidemiological connectivity among host populations due to their feeding habits, possible short visits of king penguins to conspecific colonies may not. One related key issue is whether strains of *P. multocida* against which birds have mounted an antibody response are different or similar. High heterogeneity may occur among strains of infectious agents to which hosts can be exposed at various locations, depending on evolutionary, epidemiological and ecological processes (Grenfell et al. 2004). Direct detection of DNA of the bacterium by PCR in cloacal or oral swabs could provide evidence of the local exposure to the agent. However, microbial culture of the bacterium provides more material for genetic investigations, which is critical to compare strains and potentially conduct phylogenomic studies to infer epidemiological processes. On Amsterdam Island, the isolation has been facilitated by using a chromogenic selective medium (CHROMAgar Pasteurella©) to streak swabs from organs of necropsied birds on plates. Conversely, attempts to isolate the bacterium from swabbing live birds have been much less successful, even from brown skuas on Amsterdam Island, where 98% of individuals are seropositive and >20% PCR positive (Gamble et al. 2020a, Lamb et al. 2023). At sites other than Amsterdam Island, limited exploration has yet been done by direct detection of the bacterium, but DNA of the bacterium could be detected by PCR in cloacal swabs from lesser sheathbills of Possession Island, Crozet (Bralet et al. 2025), and more broadly among other seabird species (Bralet et al. 2026), confirming their potential local exposure to the bacterium. Our serological results highlight the value of further investigation using direct detection and sequencing methods. Investigations based on genetic sequences would be useful to infer basic epidemiological parameters, such as the spatial and temporal variability in the prevalence of DNA from different strains, but also to infer transmission between locations and hosts, as well as the nature of virulence genes. In this respect, it is interesting that the recently reported strain of *P. multocida*, which was associated with a massive die-off of Adélie penguins on the Antarctic Peninsula, belongs to the same MSLT genotype as the main strain responsible for die-offs on Amsterdam Island (Ievolino et al., 2025). However, further research is required, such as the investigation of full bacterial genomes and the use of isolates from more sites, to gain a better understanding of how the bacteria circulate.

### Variability in virulence and potential for outbreaks

A large amount of knowledge is available on virulence factors in *P. multocida* in domesticated animals, notably on the role of lipopolysaccharide (LPS) (Harper & Boyce 2006). Given such knowledge, exploring the characteristics and potential diversity of such virulence factors among the strains to which seabirds are exposed could be helpful. Investigating whether pathogenic effects differ among host species and how much strains are shared would also be important. Environmental conditions at breeding colonies may also be critical to explain transmission and risks of outbreaks. It is notably expected that introduced mammal species, such as rats (Curtis 1983), and other factors, such as humidity and temperature of the colony substratum (Qin et al. 2017), could affect eco-epidemiological dynamics. The remoteness of many of the sites explains the limited availability of field data, but the results obtained for king penguins fit well with earlier reports of mortalities potentially due to avian cholera on Marion Island (Cooper et al. 2009). Also, infectious disease has been mentioned as a hypothetic cause of the dramatic decline of the once largest king penguin population on Cochon Island, Crozet (Weimerskirch et al. 2015). The results clearly warrant future investigations.

### Potential underlying immunity and selection processes

Determining whether and how antibody responses against *P. multocida* differ among species is a complementary critical issue as it may greatly affect the epidemiological interpretation that can be made from serological patterns (Pepin et al. 2017, Gamble et al. 2020b). The serological signature of exposure to the disease agent may differ depending on the temporal dynamics of the titres of antibodies after exposure and the timing of sampling (Wilber et al. 2020). Data from the longitudinal monitoring of the immunological status of individuals would be useful to complement cross-sectional surveys (Gamble et al. 2019, 2020b). Among Indian yellow-nosed albatrosses, a relatively short temporal persistence of antibody levels was detected between years on Amsterdam Island (Gamble et al. 2019). Still, repeated sampling and serological testing of individuals have yet to be conducted for other seabirds exposed to the bacterium. Studying the dynamics of antibody levels throughout the annual cycle of a resident species like the lesser sheathbill could be especially interesting to infer the role of re-exposure on the temporal antibody dynamics in such a key sentinel species. Increasing sample sizes for some species or locations may also be important, e.g., for burrow nesting petrels and South Atlantic islands.

Knowledge of the dynamics of antibody response following exposure to *P. multocida* would especially be important for interpreting results obtained in penguins. In particular, it is unclear why so low antibody prevalence levels were found in Northern rockhopper penguins on Amsterdam Island, where they are subject to local massive die-offs (Jaeger et al. 2018) and the bacterium has been isolated from dead corpses (Hashish et al. 2023 and unpublished); is their antibody response impaired or do only a few individuals mount a response because most die during an outbreak? The presence of antibodies is also important because they could play a role in the eco-epidemiological dynamics via protective effects (van Dijk et al. 2021). In anatids exposed to avian cholera, relatively short temporal persistence of antibody levels have been reported (Samuel et al. 2005), and it would also be imperative to explore why high seroprevalence and titres were found in the brown skua and giant petrels, with the potential key roles of re-exposure and properties of their humoral immune system. Combining more detailed knowledge on the exposure of seabird hosts to pathogens with demography and immunogenetics could provide important complementary insights to understand the dynamics of host-pathogen interactions in such systems. Heterogeneity in individual susceptibility to pathogens could lead to strong selection pressure throughout an outbreak in which most individuals are exposed to a pathogen. This could be the case for densely breeding species like penguins or scavengers and avian predators. In the latter case, it is interesting to note that giant petrels, skuas, or large gulls have, in several instances, been involved in short-term bursts of mortalities related to exposure to *P. multocida* (Leotta et al. 2006, Wille et al. 2016). Conversely, in an endemic situation of avian cholera like on Amsterdam Island, almost all skuas have antibodies against *P. multocida*, with no suspicious deaths for adults of that species recorded in recent years (Gamble et al. 2020a). This might be due to the early die-off of sensitive individuals, leading to the selection of resistant individuals, which can feed on chicks of other seabird species that die of avian cholera.

## Conclusions

Overall, this study highlights that carefully designed integrated approaches conducted at multiple sites on multiple species can significantly increase the understanding of the eco-epidemiological dynamics occurring in highly spatially structured colonial vertebrate communities. We showed that the agent of avian cholera *P. multocida* is broadly distributed among southern seabirds and that four scavenging and avian predator species could be efficiently used as sentinels at different spatial scales. Detecting the presence and assessing the spatial distribution of multi-host infectious agents is a prerequisite for evaluating their potential role in the dynamics of such ecosystems.

## Methods

### Study area and populations

Southern latitudes host very large populations of seabirds, a large proportion of which breed in colonies at a limited number of sites in Antarctica and on oceanic islands (Marchant & Higgins 1990). Among them, Amsterdam, Crozet, Kerguelen, and Prince Edwards Islandsin the Indian Ocean, the Tristan da Cunha Islands and the Falkland Islands in the Atlantic Ocean, and the Antarctic Peninsula, represent key biodiversity hot spots (**Figure 2**; Marchant & Higgins 1990, Ryan et al. 2009, Delord et al. 2013). In addition to marine mammals such as fur seals (*Arctocephalus* spp.) and southern elephant seals (*Mirounga leonina*), a large part of the land-based biomass is comprised of millions of breeding seabirds, notably penguins, albatrosses, petrels, shags and terns, which feed at sea, often far from their colony (Guinet et al. 1996). Moreover, a few species of scavengers and avian predators, such as giant petrels, skuas and sheathbills, are widespread and feed at colonies of marine vertebrates. Exposure of vertebrate hosts to a diversity of infectious agents has been described in those areas (Barbosa & Palacios 2008, Uhart et al. 2018), but until recently most reports concern one or a few locations or species (e.g., Leotta et al. 2006). Large outbreaks were nevertheless reported in the last years, associated with HPAI (Banyard et al. 2023, Clessin et al. 2025, McInnes et al. 2026) and avian cholera (Taylor et al. 2025, Iervolino et al. 2025).

### Sampling design for serology

Our aim was to conduct spatially extensive surveys of representative colonies of seabird species to detect evidence of exposure to *P. multocida* at a broad hierarchy of scales. To do so, we collected samples from adults of 18 species across 11 islands (**Figure 2**). Within each area, not all colonies could be sampled, but representative samples could be gathered at important series of locations (**Table 1**). For instance, on Crozet archipelago, two of the three main islands, Possession Island (46°25’S,51°45’E) and Île aux Cochons (49°28’,70°2’), were sampled, but not Île de l’Est. Within those islands, the main king penguin colony of Île aux Cochons could be sampled as well as four of the five main colonies on Possession Island (**Figure 2**). On Kerguelen archipelago (49°15′S, 69°10′E), king penguins were sampled at two large colonies, at Cap Ratmanoff and Anse de l’Antarctique, and other species could be sampled on the Courbet Peninsula, on Mayes Island in the Golf du Morbihan and on the southern-eastern tip, at Canyon des Sourcils noirs (**Figure 2**). On Amsterdam Island (37°49’S, 77°33’E), most sampling was conducted on Entrecasteaux cliffs. At the Prince Edwards Islands, samples were gathered from Marion Island (46°52’S, 37°51’E). In the Falkland Islands, samples were collected at New Island (51°43′S, 61°18′W; **Figure 2**), on the western coast of the archipelago. Samples also were collected on Gough Island (40°19′S, 9°56′W), central South Atlantic Ocean, and Saint Paul Island (38°43′S, 77°31′E), near Amsterdam Island (**Figure 2**). On the Antarctic Peninsula, samples were gathered on Byers Peninsula, on Livingston Island, South Shetland Islands (62°38′S, 61°05′W). All data presented are novel, apart for series of samples from Amsterdam Island on brown skuas and yellow-nosed albatrosses that were already published (Gamble et al. 2019, 2020a, Lamb et al. 2023).

We grouped the sampled species into five categories based on the knowledge available regarding their foraging and general ecology using a Principal Component Analysis approach (**Supplement S2**): (i) the apex predators and scavengers (skuas, giant petrels and sheathbills), (ii) the ground nesting inshore meso predators (gulls and shags), (iii) the ground nesting offshore meso predators (penguins and albatrosses) and (iv) the burrow nesting meso predators (small petrels) (**Table 1**). Each group is represented by a color for this study, respectively (i) red, (ii) blue, (iii) green and (iv) yellow (**Table 1**). Given that breeding is highly synchronous and most species are present on land in the austral summer months (October-March), sampling was conducted at that time. With respect to the years of sampling, the period covered was mostly from 2017 until 2022 (**Table 1**), with most colonies being sampled once during summer months, although some colonies, such as those of king penguins on Possession Island, Crozet, were sampled several times (2017, 2021 and 2022). Such time series enabled exploring a possible temporal stability in the pattern of antibody level distribution among species.

Individuals were captured at random on breeding, foraging or loafing sites using a pole with a noose, hands or hand nets depending on species. They were then sampled and either ringed or color marked on feathers to avoid resampling within the capture session. Blood samples of about 1 ml were taken from the ulnar or tarsal vein using heparinized 1 ml or 2.5 ml syringes. Blood samples were centrifugated in the field within a few hours of collection (c. 3000 rpm for c. 5 min). Plasma was separated from red blood cells and samples were kept in a cooler in the field until returned to the lab and stored at −20°C. When handling the birds and when circulating in the field, specific care was taken regarding biosecurity issues to limit risks of transfer of infectious agents between birds and between (sub-)colonies. This involved wearing sets of waterproof overalls that were disinfected between contacts with any two birds. Any other equipment (e.g., caliper, ruler) and hands were disinfected between birds, respectively using hydrogen peroxide, commercial disinfectant wipes, and hydroalcoholic gel. All pieces of clothing were thoroughly cleaned and disinfected again at the end of each day. On Amsterdam Island, different sets of clothing and rubber boots were used among colonies (Bourret et al. 2018). All sampled birds were in apparent good health at capture and when released after sampling.

### Serological analyses

Antibody levels against *P. multocida* in plasma samples of seabirds were measured using an immunoassay microagglutination test (MAT; SEROPAST®, Ceva Biovac, France). The assay was designed using strain ST91 according to the *P. multocida* multiple-host MLST scheme, the main strain of *P. multocida* isolated on Amsterdam Island (Jaeger et al. 2018, Bourret et al. 2018, Hashish et al. 2023). The assay had been used for quantifying the response to autologous vaccination of yellow-nosed albatross nestlings and adults, allowing to detect clear serological responses compared to controls (Bourret et al. 2018, Gamble et al. 2019). Titres above 1/10 were considered as seropositive. For brown skuas and Indian yellow-nosed albatrosses, a second immunoassay (ELISA ID Screen® *Pasteurella multocida* Chicken and Turkey Indirect) that provided results that were congruent, but less sensitive, had been used on earlier samples (Gamble et al. 2019, Gamble et al. 2020a). It was decided to use only the MAT assay, which allowed making interspecific comparisons without requiring to define a seropositivity threshold for each species (since it did not require a secondary antibody to bind to each species antibodies; Garnier et al. 2017). In order to check for the robustness of the serological assay against variability among potentially circulating strains, part of the assays were conducted in duplicate using a MAT SEROPAST® designed using a second strain of *P. multocida* isolated from an Indian yellow-nosed albatross nestling from Amsterdam Island (Hashish et al. 2023); the results showed a high congruence for both antibody titres and seroprevalence (respectively r = 0.64, p < 0.0001 and r = 0.67, p < 0.0001, n = 445 plasma samples) suggesting that the used assay is not specific to one strain and can inform on *P. multocida* exposure more broadly.

### Sexing

Most seabirds cannot be sexed visually in the field, thus we proceeded with genetic sexing (**Supplement S3**). Both species of giant petrels present a marked sexual dimorphism in size (Carlos & Voisin, 2008), thus sexing could be made from field data (male, bill length > 105mm; female bill length < 95mm), and confirmed genetically for some individuals captured in 2021 and 2022. Although for these 2 species it could be predicted that males may be more exposed to avian cholera due to sexual segregation in foraging habits, with males more foraging on land on carcasses (Gonzalez-Solis et al. 2000, Thiers et al. 2014), highly unbalanced sampling towards males due to our sampling efforts on penguin colonies prevented us from meaningfully testing this prediction.

### Statistical analyses and mapping

Mean titres (+- SEM) and proportions of individuals that showed titre values above the 1/10 threshold were used to make comparisons among species groups and species/site/year/sex for relevant subsets. Generalized linear mixed models (with Poisson and binary distributions) were implemented in R (package ‘ggeffect’, Lüdecke 2018) to test for mean titres and seropositivity as a function of combined associations of species, site and year, on appropriate subsamples. When possible, given the available data, 2-way interactions were considered (e.g., to test for a potential interaction between site and year on the proportion of seropositive king penguins on Possession Island, Crozet, where three colonies were sampled in 2017, 2021 and 2022). Seroprevalences were calculated using package ‘prevalence’ in R (Devleesschauwer et al. 2022) to account for some possible uncertainty in the assay (for this, a sensitivity of 0.95 and specificity of 0.95 were set) and those seroprevalences were reported +-Confidence Intervals; **Table 1**). All maps were created using ‘leafletR’ package (Graul & Christian 2016) with CartoDB.Positron as background.

## Acknowledgements

We thank Nicolas Keck, Romain Dedet, Taya Forde, Karin Lemberger (Vetdiagnostics), Corisande Abiven, Natacha Garcin, Nicolas Giraud, Marine Bely and Louisiane Burkart for help at various stages of the work and Norman Ratcliffe, Jaimie Cleeland, Kate Lawrence, Michelle Risi, Christopher Jones, Alexis Osborne and Steffen Oppel for assistance in the field. We also thank Céline Bocquet, Elodie Paciello, Nolwenn Trividic, Lucia Llorente for contributing to the field surveillance and sampling on Crozet, and Nelly Lesceau and her team (Ceva Biovac, France) for their technical assistance with the microagglutination test. We acknowledge support from the French Polar Institute (IPEV ECOPATH-1151, ORNITHOECO-109, OISEAUX PLONGEURS-394 and ECONERGY-119), ANR ECOPATHS (ANR-21-CE35-0016) project, BiodivRestore REMOVE_DISEASE project (ANR-21-BIRE-0006), France relance TVACALBA project, Réserve Naturelle Nationale des Terres Australes Françaises, Zone Atelier Antarctique et Terres Australes (ZATA) and OSU OREME (ECOPOP). This research was funded through the 2020-2021 Biodiversa+ and Water JPI joint call for projects under the BiodivRestore ERA-NET Cofund (GA N°101003777). This study is part of the long-term Studies in Ecology and Evolution (SEE-Life) program of CNRS and is a contribution to the Plan National d’Actions Albatros d’Amsterdam. For work in the Falklands, we thank the South Atlantic Environmental Research Institute, Falkland Islands Government, FCT (Fundação para a Ciência e a Tecnologia) – Portugal through the strategic projects UIDB/04292/2020 (MARE), LA/P/0069/2020 (ARNET) and DivRestore/0012/2020 (REMOVE_DISEASE project). The Tristan da Cunha Conservation Department and Tristan da Cunha government granted permission (Environmental Research Permit 1742) to work on the islands, and the RSPB Ethics Advisory Committee approved the sampling design. AG received a PhD fellowship from the French Ministry of Research, and JT received support from Ceva Biovac and ANRT for a CIFRE PhD fellowship. Support is also acknowledged from the Royal Society (Newton International Fellowship NIF\R1\211869; AG), the Biodiversity Challenge Fund (Darwin Plus Grant DPLUS167; AG) and Falklands Conservation. The experimental design in Terres Australes et Antarctiques Françaises was approved by French Ministry of Research (APAFIS #10257-2018011712301381v6, #31773-2019112519421390v4), Comité de l’Environnement Polaire and CNPN. The project in the Falkland Islands was approved by FIG (Research Licence R27/2017).

